# An active RNA transport mechanism into plant vacuoles

**DOI:** 10.1101/2021.07.28.454214

**Authors:** BE Floyd, MM Wong, AY Liu, SC Morriss, Y Mugume, Z Kazibwe, V Ridout, X Luo, GC MacIntosh, DC Bassham

## Abstract

RNA degradation inside the plant vacuole by the ribonuclease RNS2 is essential for maintaining nucleotide concentrations and cellular homeostasis via the nucleotide salvage pathway. However, the mechanisms by which RNA is transported into the vacuole are not well understood. While selective macroautophagy may contribute to this transport, macroautophagy-independent transport pathways also exist. Here we demonstrate a mechanism for direct RNA transport into vacuoles that is active in purified vacuoles and is ATP hydrolysis-dependent. We identify the RNA helicase SKI2 as a factor required for this transport pathway, as *ski2* mutant vacuoles are defective in transport. *ski2* mutants have an increased autophagy phenotype that can be rescued by exogenous addition of inosine, consistent with a function in nucleotide salvage. This newly-described transport mechanism is therefore critical for RNA degradation, recycling and cytoplasmic nucleotide homeostasis.

## Introduction

Nucleotide salvage, the recycling of nucleosides and nucleobases to produce nucleotides, is necessary to maintain cellular homeostasis in eukaryotes ^1–3^. Cells have the ability to produce nucleotides through de novo synthesis or through the salvage pathway, but when nucleosides or nucleobases are available, the salvage pathway is preferred due to its favorable energy balance ^4^. However, the salvage pathway is not simply a complement to the de novo biosynthetic pathway. The salvage pathway removes bases and nucleosides that can have an inhibitory effect on metabolism, and in plants it is also linked to production of cytokinins ^1^.

The main source of nucleosides for nucleotide salvage is thought to be RNA degradation ^5^, and given its abundance, ribosomal RNA is likely the most prevalent source. Previous work showed that rRNA is degraded by RNS2, the main ribonuclease in the Arabidopsis vacuole ^6^. Mutant plants lacking RNS2 activity, or plants expressing a mutant version of the protein that is mislocalized outside the vacuole, accumulate rRNA ^6,7^. These mutants also have constitutive activation of autophagy, a process usually only highly active during stress ^6–8^. Transcriptional and metabolic analyses suggested that plants lacking vacuolar RNA turnover compensate by activation of the pentose-phosphate pathway, which provides ribose-5-P to the de novo purine biosynthesis pathway ^9^. Moreover, supplementing *rns2* mutant plants with purine nucleosides (inosine or hypoxanthine) suppresses their constitutive autophagy phenotype, reinforcing the idea that defective RNA recycling leads to a decrease in purine availability that triggers autophagy as a compensatory mechanism ^10^. Genetic analyses indicated that some components of the autophagy machinery are necessary for efficient transport of rRNA ^7^ and other RNA species ^11^ from the cytoplasm to the vacuole, suggesting that a selective autophagy mechanism similar to yeast ribophagy ^12^ may exist in plants. However, rRNA transport into the vacuole was unaffected in *rns2-2 atg9-4* mutants, indicating that a macroautophagy-independent transport mechanism may also participate in nucleic acid import into the Arabidopsis vacuole ^7^.

An alternative pathway for RNA transport into lytic organelles, called RNautophagy, has been described in animal cells ^13^. In RNautophagy, RNA is transported into lysosomes by a direct import mechanism that requires at least two lysosomal proteins, Lysosomal associated membrane protein 2 isoform C (LAMP2C) and SID1 transmembrane family member 2 (SIDT2) ^13,14^; however, these proteins have not been described in plants. This transport mechanism is capable of transporting rRNA and requires ATP ^13^ but not lysosome acidification ^15^. RNautophagy seems to have a significant contribution to the degradation of cellular RNAs ^13,14^. We hypothesized that an analogous RNA transport mechanism could be present in Arabidopsis and mediate the macroautophagy-independent import of rRNA into vacuoles observed in autophagy defective mutants.

In this study, we developed assays to measure the transport of rRNA and a synthetic fluorescent RNA into isolated vacuoles. Using these methods, we demonstrated the presence in Arabidopsis of an ATP-dependent novel transport mechanism that can carry RNA into the vacuole. We also identified an RNA helicase, SKI2, that is necessary for this vacuolar transport mechanism. Our results demonstrate that plants have a mechanism analogous to RNautophagy, but requiring distinct machinery, and provide tools for the dissection of this novel mechanism that can contribute to nucleotide salvage and cellular homeostasis.

## Results

### Isolated Arabidopsis vacuoles can import exogenous rRNA

A Blastp search ^16^ of the predicted proteomes of Arabidopsis and 63 other plant species in the Phytozome database ^17^ using human and mouse SIDT2 and LAMP2C as queries did not return any significant hits. If RNA can be transported directly across the vacuolar membrane in plants, it must therefore occur via an unrelated mechanism. To test whether an autonomous vacuolar transport mechanism exists in Arabidopsis, independent of SIDT2 and LAMP2C-like proteins, we set up an in vitro system to measure direct rRNA uptake using isolated vacuoles. A challenge with quantification of RNA incorporation into vacuoles is the presence of RNS2, the main vacuolar RNase in Arabidopsis ^6,8^, which would degrade any RNA that is imported into the organelle. To avoid this problem, we isolated vacuoles from *rns2-2* null mutant plants, which accumulate RNA due to reduced degradation ^6,7^. To distinguish between Arabidopsis RNA already present in *rns2-2* vacuoles and RNA transported into the vacuole during the assay, we used Drosophila RNA as a transport substrate instead of plant RNA. A schematic of the experimental design is shown in Figure 1A. Purified vacuoles were incubated with 5 μg total Drosophila RNA in either the presence or absence of ATP. The samples were gently mixed and incubated for 5 minutes at room temperature to allow transport. To prevent contamination of minus ATP samples with cellular ATP released during vacuole purification, apyrase was added to the samples without exogenously added ATP. After allowing transport to occur, RNase A was added for 20 minutes on ice to remove RNA not incorporated into vacuoles; Drosophila RNA taken up by vacuoles would be protected from RNase A degradation. Following addition of an RNase A inhibitor, RNA was extracted from the vacuoles and quantified by qRT-PCR using Drosophila rRNA specific primers (Figure S1). Vacuolar acid phosphatase activity was used for normalization as in Floyd et al. ^7^. Using this assay, we were able to detect Drosophila rRNA uptake into plant vacuoles. Vacuoles incubated with ATP accumulated ∼3-fold more 18S and 28S Drosophila rRNA than those without ATP (Figure 1B). Control samples lacking exogenous RNA showed no accumulation of rRNA, indicating that the Drosophila rRNA primers can discriminate between Drosophila rRNA and the Arabidopsis rRNA already present in the vacuole. Control samples lacking purified vacuoles show the quantity of detectable RNA remaining after RNase A treatment (Figure 1B). These results indicated that vacuoles purified from *rns2-2* plants can import RNA in an ATP-dependent manner, analogous to the RNautophagy mechanism reported in mammals ^13^.

**Figure 1.**
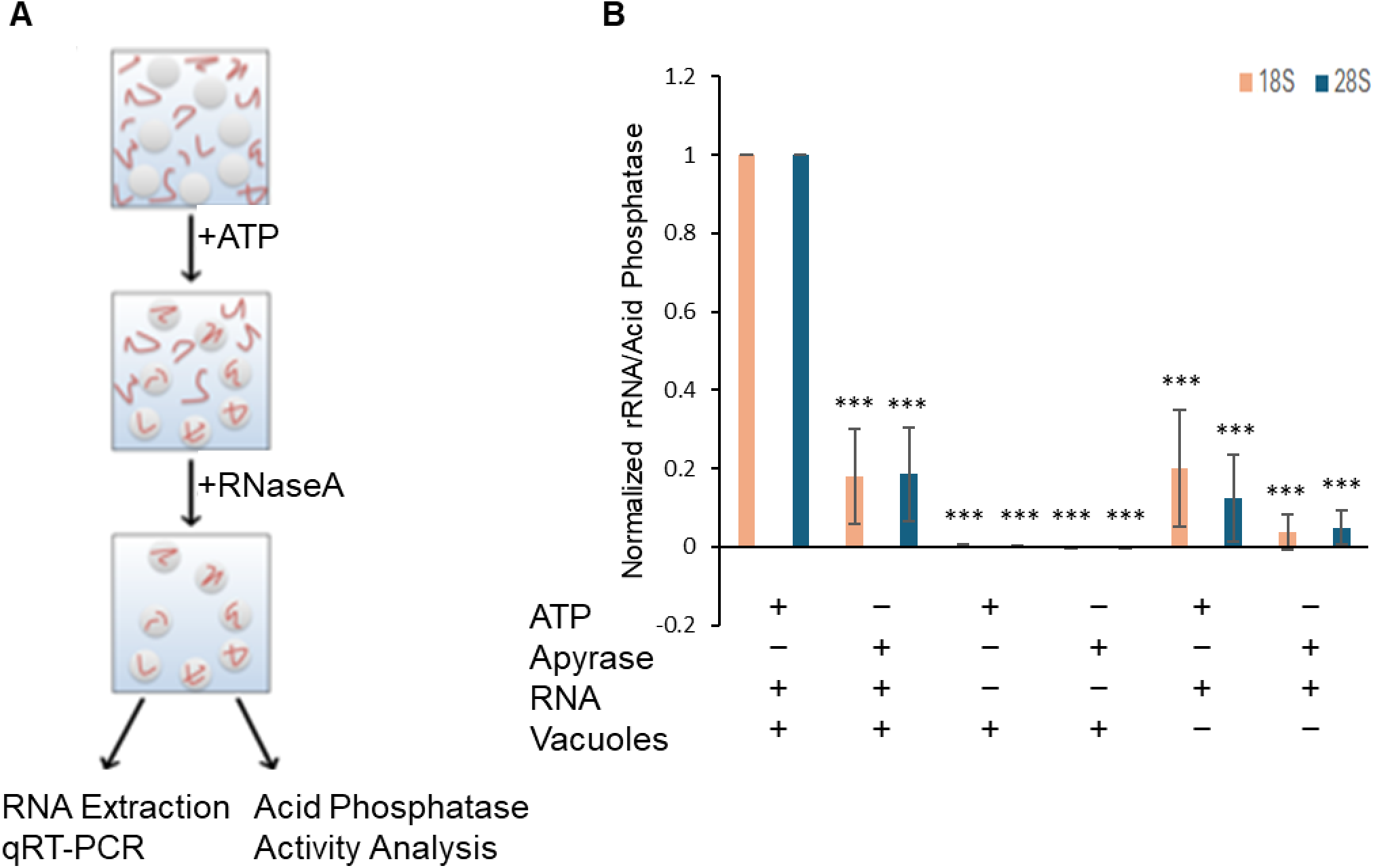
RNA is transported into plant vacuoles in a process that requires ATP. (**A**) Schematic of the experimental design to measure the transport of Drosophila RNA (squiggles) into purified Arabidopsis vacuoles (circles). Vacuoles were incubated with RNA in the presence or absence of ATP. After incubation, unincorporated RNA was degraded with RNase A. RNA imported into vacuoles was protected from degradation, and was quantified using qRT-PCR. Vacuolar acid phosphatase activity was used for normalization. (**B**) Quantification of RNA transport into *rns2-2* vacuoles. RNA transport was analyzed with or without addition of ATP. Apyrase was used to remove residual ATP in the –ATP treatment. Abundance of 18S and 28S Drosophila rRNA incorporated into vacuoles was quantified by qRT-PCR, and the results were normalized to acid phosphatase activity. Samples were analyzed in triplicate for three biological replicates, separately for 18S and 28S rRNA, and normalized to vacuolar rRNA transport with addition of ATP. Error bars represent standard deviation. Asterisks indicate significant difference according to one-sample t test (*** *p* ≤ 0.001) with respect to vacuolar rRNA transport with addition of ATP.

### Fluorescent synthetic RNA can be used to assay vacuolar trans-membrane transport activity

To confirm the results of RNA transport identified by qRT-PCR analysis, an assay was developed to more directly visualize RNA transport into purified vacuoles using fluorescent RNA (FL-RNA). We used a commercially synthesized 15-nucleotide random RNA sequence conjugated at the 3’-end with Alexa488 fluorophore as transport cargo. A schematic of the experimental design is shown in Figure 2A. Wild type vacuoles were used in this analysis since intravacuolar RNA degradation should not interfere with the quantification of fluorescence inside the vacuole. Purified vacuoles were incubated for 20 minutes with FL-RNA in the presence of ATP to promote transport; room temperature incubation was found to be unnecessary as measurable transport occurred on ice, under which conditions stability of the vacuoles is improved. Following FL-RNA transport incubation, samples were diluted ten-fold with vacuole buffer to dilute out non-imported FL-RNA, reducing background fluorescence. Vacuoles were then imaged using microscopy. Using this assay, we were able to observe fluorescence accumulation inside isolated Arabidopsis vacuoles (Figure 2 B-D). We observed three classes of vacuole, exhibiting either zero, low or high fluorescence (Figure 2B).

**Figure 2.**
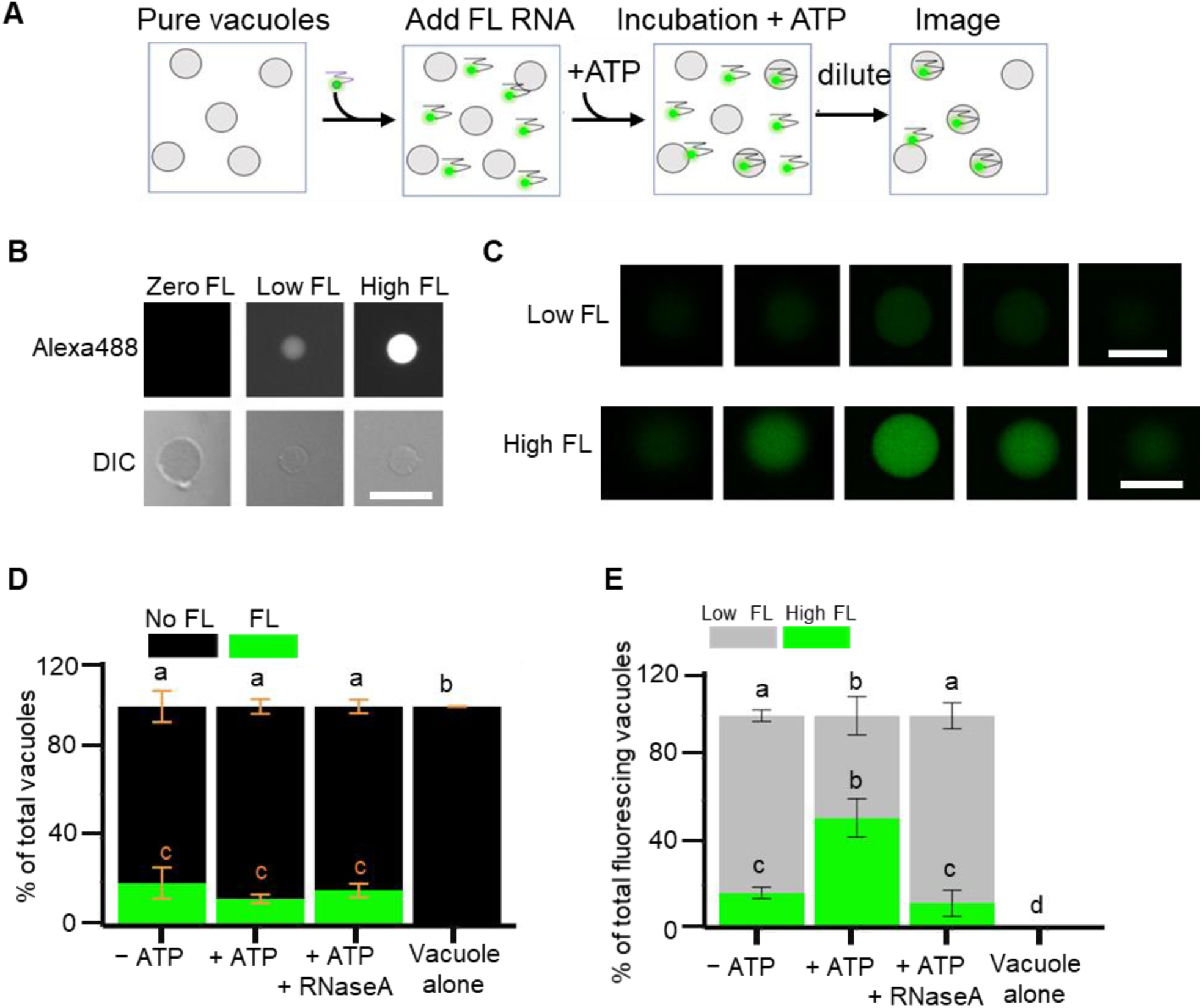
Alexa488-tagged-RNA is transported into plant vacuoles in vitro. (**A**) Schematic of experimental design for transport of fluorescently labeled RNA (in green) to purified vacuoles (circles). Vacuoles were incubated with FL-RNA in the presence or absence of ATP, diluted to reduce background fluorescence, and incorporation of RNA into vacuoles was determined by microscopy. (**B**) Fluorescence microscopy and differential interference contrast (DIC) microscopy images of vacuole examples with zero, low, and high fluorescence (FL). Scalebar = 25μm. (**C**) Confocal microscopy z-series of 2.7 μm optical sections of low (top panel) and high (bottom panel) fluorescing vacuoles. Scalebar = 20μm. (**D**) Quantification of the number of fluorescent vacuoles for three biological replicates normalized to the total number of vacuoles. (**E**) Fraction of low and high fluorescence vacuoles normalized to total number of fluorescent vacuoles. Error bars represent ± SD. Similar letters indicate no significant difference according to one-way analysis of variance (ANOVA) with Tukey’s multiple comparisons test. For (D), a, b; b, c; a, c p < 0.0001. For (E) a, b; b, c; b, e p < 0.01, a, c; a, e p < 0.0001, c, e p < 0.05.

To determine if the observed fluorescence was inside the vacuole or binding to the outer surface, vacuoles were imaged using confocal microscopy. Vacuoles with either low or high fluorescence were imaged from multiple focal z-layers and for both high and low fluorescence vacuoles fluorescence was found throughout the organelle at multiple z-positions rather than the outer edges alone (Figure 2C). The fluorescent RNA is therefore within the lumen of the vacuole rather than bound to the membrane. These observations indicate that FL-RNA can be used to visualize the RNautophagy-like transport pathway in purified Arabidopsis vacuoles, and that the transport mechanism can transport small RNAs in addition to rRNA.

### The efficient transport of FL-RNA into vacuoles needs ATP hydrolysis

After determining that the observed fluorescence was originating from the vacuole interior for both low and high fluorescence vacuoles, we assessed the requirements for transport into the isolated wild-type vacuoles. First, we determined the percentage of vacuoles observed to be fluorescent in the presence or absence of ATP. About 20% of vacuoles showed a fluorescence signal regardless of treatment (Figure 2D), with the remainder being non-fluorescent. Vacuoles alone, without addition of FL-RNA, showed no autofluorescence. This suggests that only a subpopulation of vacuoles is competent for transport, although the reason for this is unknown.

We then determined the distribution of vacuole fluorescence intensities in the FL-RNA transport system. Vacuoles were manually assigned to different categories, high FL, low FL, and zero FL (Figure 2B and 2E). Vacuoles showing a fluorescence intensity that saturated or nearly saturated the detector were assigned to the high-FL group. Vacuoles showing all other amounts of fluorescence were assigned to the low-FL group. Vacuoles showing no fluorescence were assigned to the zero-FL group. Using this ranking for quantification, there was a significant increase in vacuoles in the high FL population when ATP was added (Figure 2E), suggesting that efficient transport requires ATP. We confirmed that the observed transport of the fluorophore into the vacuoles required the presence of the RNA, and that the Alexa488 fluorophore alone was unable to enter the vacuoles by a non-selective process. For this, the FL-RNA probe was incubated with RNase A to digest the RNA portion of the probe prior to incubation with vacuoles and ATP. RNase A treatment of the FL-RNA probe resulted in few high-FL vacuoles, indicating that RNA enhances transport of the Alexa488 fluorophore (Figure 2E). The number of low FL vacuoles was similar in the minus ATP samples and in the samples with probe pre-treated with RNase A (Figure 2D and 2E). These results suggest that the low FL vacuoles represent non-selective transport that is independent of RNA and ATP, while high FL vacuoles represent an ATP-dependent RNA transport system. Identical results were obtained using vacuoles purified from *rns2-2* plants (Figure S2), indicating that the same RNA-transport mechanism is functional in the wild-type and mutant backgrounds.

To eliminate bias in our analysis, we used Fiji software ^18^ to measure the fluorescence intensity of individual fluorescing vacuoles. A circular region within the vacuole being measured was quantified, and average intensities for each sample were statistically compared (Figure 3). Wild type vacuoles treated with ATP showed the highest fluorescence intensity, as well as the widest distribution of fluorescence intensities. Removal of ATP resulted in overall decreased fluorescence and a narrower distribution (Figure 3A). Although bright vacuoles were seen in minus ATP and in the RNase A-treated probe samples, they were mostly statistical outliers following analysis (vacuole FL > 1.5 standard deviations away from sample mean). Background fluorescence was quantified to illustrate the level of fluorescence from non-transported FL-RNA remaining after sample dilution. Vacuoles alone showed no quantifiable autofluorescence. Similar results were obtained for *rns2-2* vacuoles (Figure 3B).

**Figure 3.**
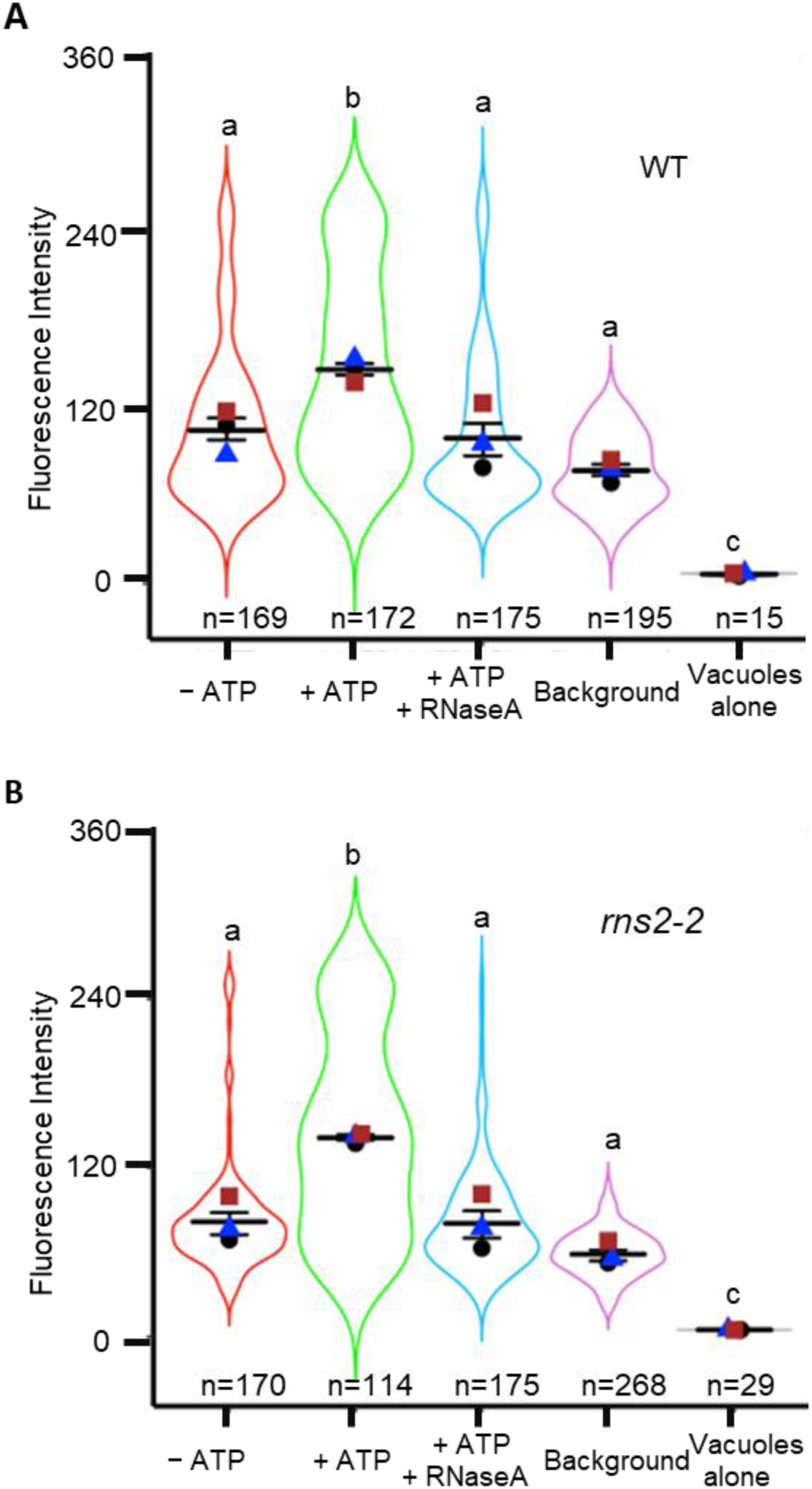
High fluorescence vacuoles are increased in the presence of ATP. FL-RNA transport was analyzed using vacuoles purified from wild type (**A**) or *rns2-2* mutant (**B**) plants. Fluorescence intensity for each vacuole was determined from microscopy images using Fiji software. Each experiment was repeated three times. The number of vacuoles analyzed is indicated below each plot. Larger horizontal bar indicates mean, error bars represent ± SD. The mean and SD were computed from the means of three biological replicates, indicated by blue triangle, brown square and black circle. Similar letters indicate no significant difference according to one-way ANOVA followed by post-hoc pairwise Student’s *t*-test. For (A) and (B), a, b *p* < 0.05; b, c *p* < 0.001; a, c *p* < 0.0001.

To further elucidate the requirements of the vacuolar RNA transport mechanism, we tested the effect of inhibitors on FL-RNA transport. We first tested whether ATP hydrolysis was required or whether ATP binding alone was sufficient for RNA transport. The addition of adenosine 5’-(β, γ-imido) triphosphate to the vacuole transport assay, which contains a non-hydrolysable bond between the β and γ phosphates, significantly reduced the mean vacuole fluorescence and lowered the fluorescence distribution (Figure 4A). This indicates that ATP hydrolysis was required for efficient transport. A common ATP-dependent mechanism of trans-membrane transport in many organisms involves ATP-binding cassette (ABC) transporter proteins ^19^.

**Figure 4.**
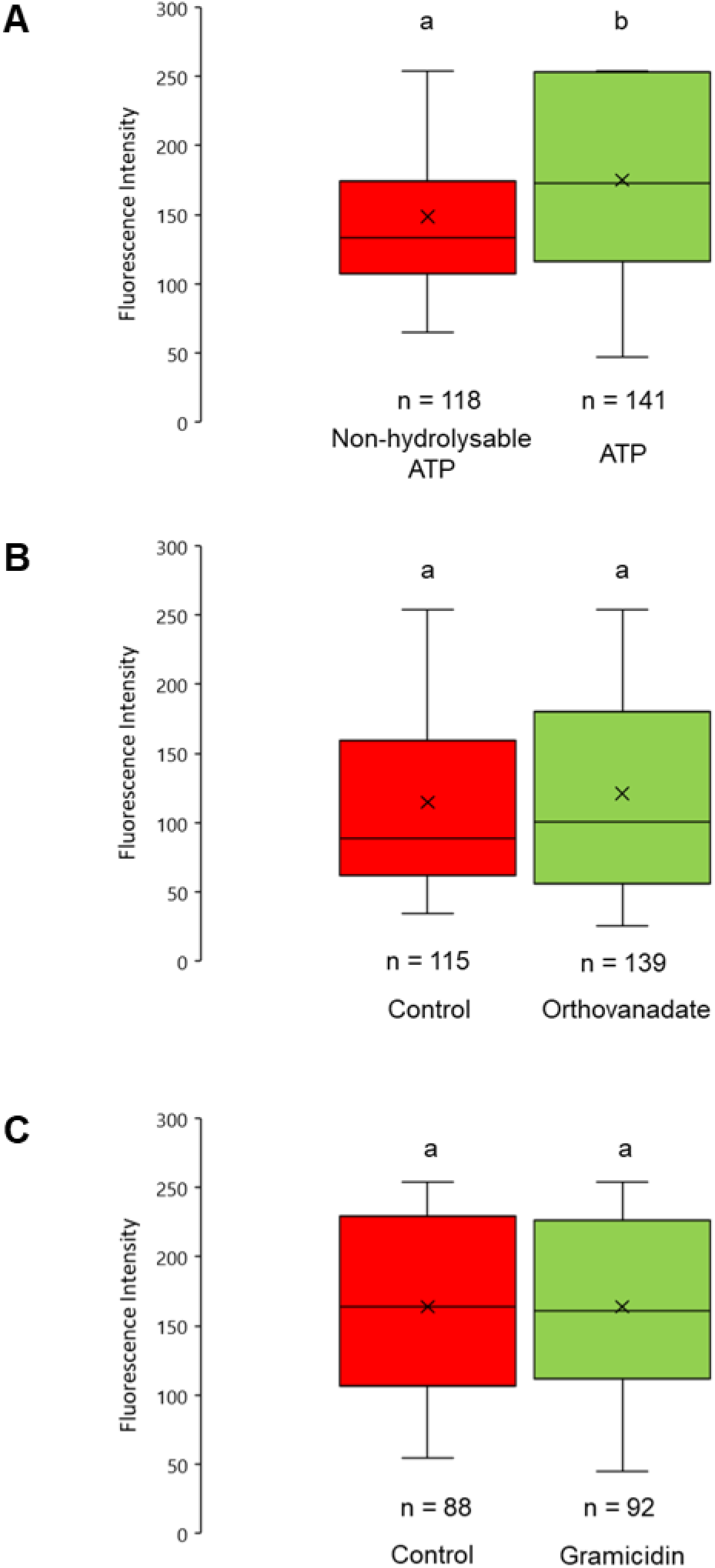
ATP hydrolysis is necessary for efficient RNA transport into vacuoles. FL-RNA import into isolated vacuoles was determined by fluorescence quantification of microscopy images. (**A**) Treatment of vacuoles with a non-hydrolysable ATP (adenosine 5’-(β,γ-imido) triphosphate lithium salt hydrate). (**B**) Treatment of vacuoles with the ABC-transporter inhibitor sodium orthovanadate. (**C**) Treatment of vacuoles with the ion gradient uncoupler gramicidin. Horizontal bar indicates mean, error bars represent ± SD. The number of vacuoles analyzed is indicated below each plot. Similar letters indicate no significant difference according to pairwise Student’s two-sided *t*-test, a, b *p* < 0.05.

Treatment of vacuoles with the ABC transporter inhibitor sodium orthovanadate, a suppressor of ATPase activity ^20^, had no significant effect on FL-RNA transport efficiency (Figure 4B), suggesting that RNA transport into the vacuole does not involve an ABC transporter. Co-transporters are another class of proteins used in the trans-membrane transport of molecules into the vacuole ^21^. Plant vacuoles maintain a high concentration of ions in their lumen that drive antiport mechanisms ^21^. To test whether FL-RNA transport relies on the ion gradient in plant vacuoles, we treated vacuoles with gramicidin to depolarize the membrane ionic gradient ^22,23^. Treatment of vacuoles with gramicidin had no significant effect on FL-RNA transport efficiency (Figure 4C), indicating that RNA transport into the vacuole does not require an ion gradient to drive transport.

### SKI2 is necessary for FL-RNA transport into vacuoles

As a first attempt to identify proteins involved in this vacuolar RNA transport mechanism, we used a candidate gene approach. We searched the published tonoplast proteome of Arabidopsis vacuoles ^24^ for proteins with RNA-binding motifs. In this proteome dataset we identified two proteins annotated as RNA helicases (encoded by *At3g06480* and *At3g46960*) as the top candidates, as they possessed RNA-binding domains and ATP hydrolysis activity. *At3g46960* corresponds to *SUPERKILLER 2* (*AtSKI2*), the gene encoding a DExD⁄H box RNA helicase associated with the cytoplasmic RNA exosome ^25,26^; while *At3g06480* corresponds to *RNA Helicase 40* (*RH40*), encoding a DEAD box RNA helicase with homology to helicases involved in rRNA processing ^27,28^. To determine if either of these proteins had a role in the vacuolar RNA transport mechanism, we obtained T-DNA insertion Arabidopsis lines (termed *ski2-5*, *rh40-1* and *rh40-2*) that disrupted the corresponding genes and identified individual plants homozygous for each mutation, which lacked expression of the corresponding target gene (Figure S3). Then, we isolated vacuoles from the *ski2* and *rh40* mutant lines and tested their ability to transport RNA using the FL-RNA system.

Vacuoles isolated from the *ski2-5* mutant plants had significantly reduced FL-RNA transport capability, and this vacuolar RNA transport was restored to wild-type levels after complementation of the *ski2* mutant with an extra copy of the *SKI2* gene (Figure 5A and Figure S3). On the other hand, vacuoles from two independent *rh40* mutants had normal FL-RNA transport levels (Figure 5B). These results indicated that SKI2 is necessary for the Arabidopsis RNautophagy-like mechanism, while RH40 is not.

**Figure 5.**
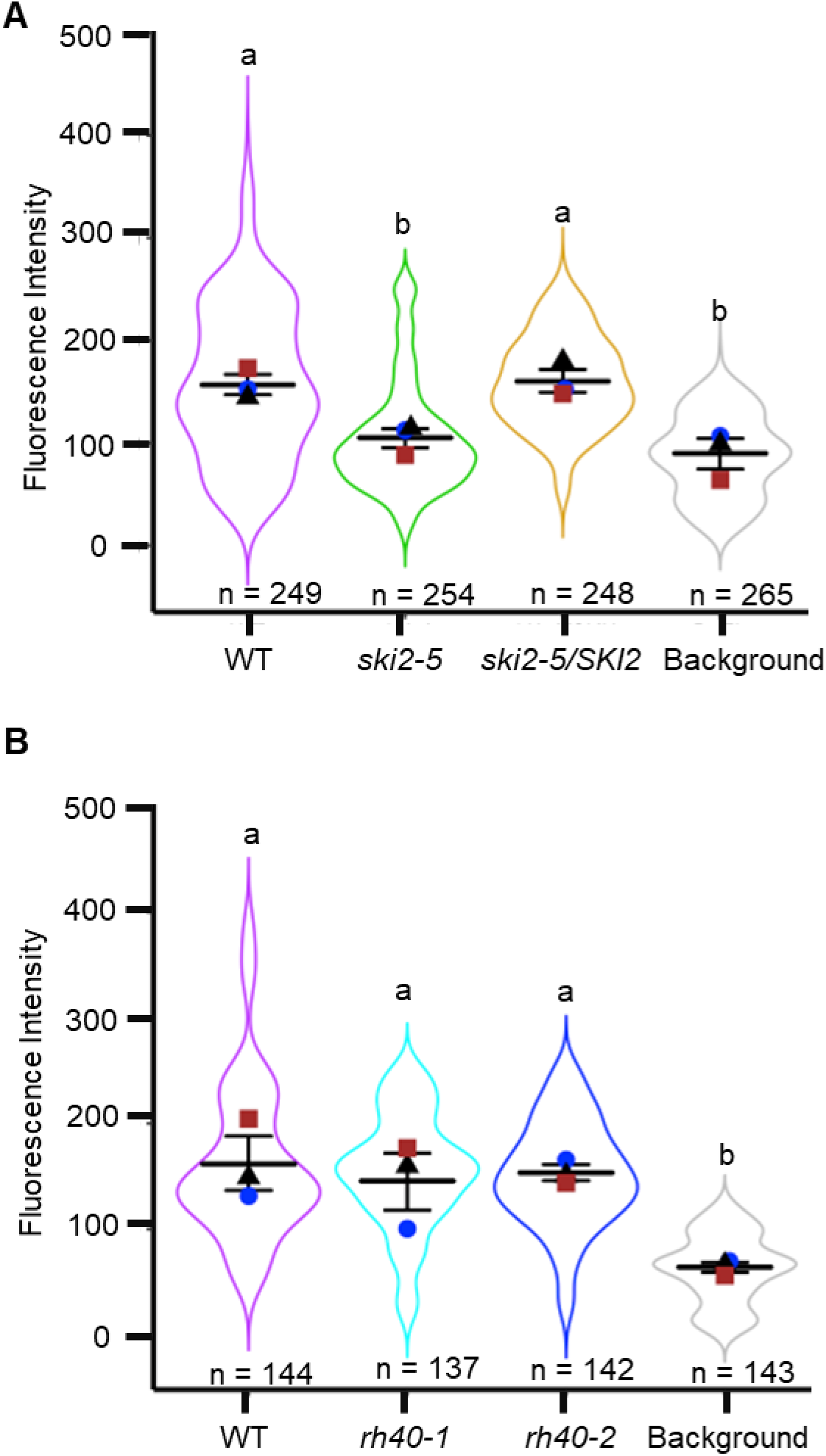
SKI2 is necessary for efficient RNA transport into vacuoles. (**A**) FL-RNA transport into vacuoles isolated from wild type, *ski2-5*, and the *ski2-5* line complemented with a wild type copy of the *SKI2* gene (*ski2-5*/*SKI2*). (**B**) FL-RNA transport into vacuoles isolated from wild type and the *rh40-1* and *rh40-2* lines. The total number of vacuoles is indicated below each plot. Larger horizontal bar indicates mean, error bars represent ± SD. The mean and SD were computed from the means of three biological replicates, indicated by blue circle, brown square and black triangle. Similar letters indicate no significant difference according to one-way ANOVA followed by post-hoc pairwise Student’s *t*-test; a, b *p* < 0.05.

Vacuolar RNA degradation is critical for the maintenance of nucleotide homeostasis ^9,10^. Mutants with deficiencies in the nucleotide salvage pathway, including those with defects in vacuolar RNA degradation such as *rns2* ^6–8^, have constitutive activation of the autophagy pathway. We therefore tested whether the *ski2-5* mutant has constitutive autophagy, indicating a physiological role in maintaining nucleotide concentrations. An increase in flux through the autophagy pathway in *ski2-5* mutant seedlings was indicated by immunoblotting with ATG8 antibodies in the presence or absence of concanamycin A to block vacuolar degradation. As ATG8 is degraded during autophagy, the accumulation of ATG8 protein is a proxy for the activity of the autophagy pathway ^29^. An increase in ATG8 was seen in the *ski2-5* mutant, and this increase was rescued in the complemented line (Figure 6A, B).

**Figure 6.**
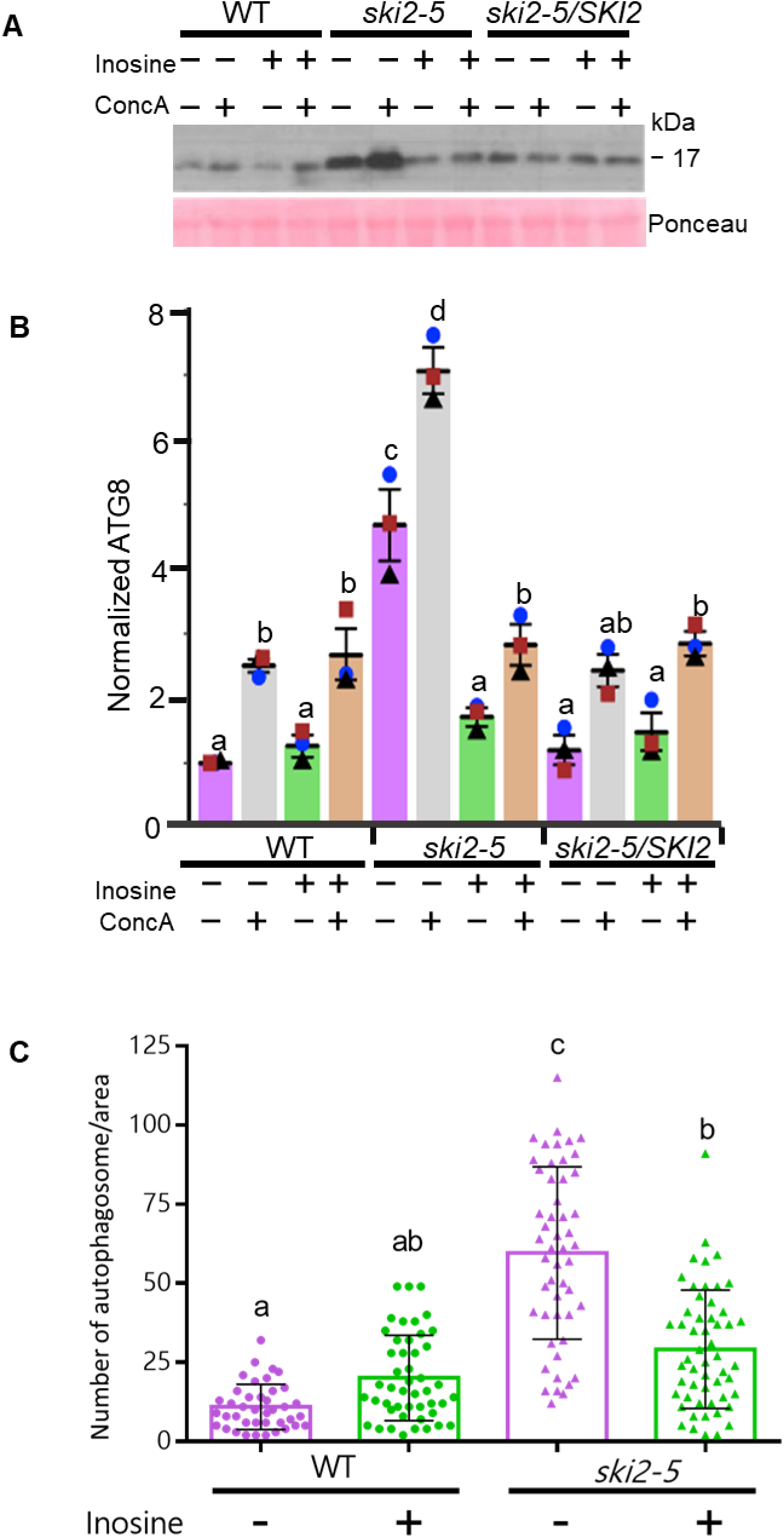
*SKI2* deficiency causes increased basal autophagy that is reduced by inosine treatment. (**A**) Seven-day old seedlings of the indicated genotypes were incubated with or without 50 µM inosine (final concentration) in liquid ½ MS medium supplemented with or without 1 µM final concentration of concanamycin A (ConcA) for 12 hours, and total protein was separated by SDS–PAGE followed by immunoblotting with anti-ATG8 antibodies. Shown are representative images from three independent biological replicates. (**B**) Quantification of the ATG8 band intensity, compared to the Ponceau S-stained band corresponding to Rubisco large subunit, and normalized using the wild type value. Larger horizontal bar indicates mean, error bars represent ± SD for three biological replicates (blue circle, black rectangle and brown square). Similar letters indicate no significant difference according to pairwise Student’s two-sided equal variance *t*-test. a, b *p* < 0.05; a, c *p* < 0.001; a, d *p* < 0.0001; b, c *p* < 0.05; b, d *p* < 0.01; c, d *p* < 0.05. (**C**) Seven-day-old seedlings of the indicated genotypes expressing the GFP-ATG8e autophagosome marker were imaged by confocal microscopy and autophagosomes counted per unit area for three biological replicates. Similar letters indicate no significant difference according to one-way ANOVA with Tukey’s multiple comparisons test using Graphpad Prism. In all cases, *p* < 0.05.

Constitutive autophagy has been observed for *rns2* mutants, which are defective in the vacuolar RNA salvage pathway, but disturbance in other pathways that affect cellular homeostasis can also trigger an autophagy phenotype ^30^. However, the increase in autophagy observed in *rns2* lines can be reversed by incubation in media containing purines, supporting the role of the RNase in nucleotide homeostasis maintenance ^10^. Thus, to test whether SKI2 participates in the RNA salvage pathway, we used inosine treatments to chemically complement the cellular phenotype. The addition of this purine to the medium was able to reverse the *ski2-5* autophagy phenotype, consistent with a role in the RNA salvage pathway (Figure 6). To confirm these results, the autophagosome marker GFP-ATG8e was introduced into the *ski2-5* mutant and autophagosome numbers counted. Compared with WT seedlings, *ski2-5* mutant seedlings had increased numbers of autophagosomes, and this increase was suppressed by inosine addition (Figure 6C). Together these results support the hypothesis that *ski2-5* functions in vacuolar RNA uptake as a component of the vacuolar RNA salvage pathway.

## Discussion

The vacuole RNA salvage pathway is an important component of nucleotide homeostasis in plants ^31^. While the main vacuolar RNase involved in this process has been well-characterized ^6,8,32^, the mechanisms that translocate RNA from the cytoplasm to the vacuole are less understood. Here, we established a straightforward assay to study RNA transport and identified a novel mechanism present in Arabidopsis vacuoles. This vacuolar RNA import system seems functionally analogous to the RNautophagy mechanism described in animals. Animal RNautophagy can transport rRNA and short oligonucleotides ^13,33^ through an ATP-dependent process that functions in isolated lysosomes ^13,15^. We found that, in a similar fashion, isolated vacuoles can transport rRNA and short oligonucleotides in an ATP-dependent manner.

Moreover, we showed that ATP hydrolysis is necessary for optimal transport and that neither ABC transporters nor a proton gradient are involved in this process. Interestingly, mutant mice defective in the RNautophagy process have an induced muscular dystrophy-like phenotype, and skeletal muscle fiber cells show an increase in autophagolysosomes ^34^, and Arabidopsis mutants defective in vacuolar RNA transport show an increase in autophagy. However, the plant and animal systems cannot be mediated by the same mechanism. Animal RNautophagy requires two lysosomal proteins, LAMP2C and SIDT2 ^13,14^. These proteins function synergistically to bind and transport RNA ^35^. Both proteins localize to the lysosome membrane ^14,36^ and contain arginine-rich RNA binding motives in their cytosolic domains and show preference for RNA with poly(G) sequences ^33,35,37^. However, genes encoding LAMP2C and SIDT2 are absent from plant genomes ^31^, and therefore the pathway we have discovered in Arabidopsis must use a distinct protein machinery. Thus, the plant vacuolar RNA transport uses a different, previously undescribed, mechanism for RNA import into the lytic organelle. While it has been proposed that the animal RNautophagy involves transport through a channel-type mechanism, we cannot discard alternative mechanisms in plants. For example, the transport observed in our experiments could be a type of microautophagy, a process that uses different mechanisms involving invagination of the vacuole membrane ^38^. Although less well-characterized, the presence of microautophagy processes in plants has been reported ^39–41^.

Our candidate gene approach identified SKI2 as a putative component of the transport system. Vacuoles from *ski2* mutant plants have compromised RNA transport and show the same constitutive autophagy phenotype as *rns2* mutants that lack the main vacuolar RNase activity ^6,7^. Moreover, altered nucleotide homeostasis in mutants with defective vacuolar RNA salvage can be compensated by feeding seedlings with purine nucleosides, and the *rns2* constitutive autophagy phenotype is chemically complemented by inosine feeding in a dose-dependent manner ^10^. Consistent with a role in the RNA salvage pathway, the *ski2* autophagy phenotype is also complemented by inosine feeding and SKI2 may contribute to nucleotide homeostasis through both the vacuolar and cytoplasmic RNA salvage pathways. We show here that SKI2 is necessary for the RNautophagy-like mechanism that delivers RNA to the vacuole for degradation. In addition, as part of the RNA exosome ^25,26^, SKI2 mediates several processes that involve cytoplasmic degradation of cellular RNAs, including normal mRNA decay ^26,42^, and quality control mechanisms such as nonsense mediate decay ^43,44^, no-go decay ^45^, nonstop decay ^46^, and nonfunctional rRNA decay ^47^. A role in nucleotide homeostasis for RNA exosome-mediated RNA decay has also been proposed in animals ^48^.

The detection of SKI2 in the vacuole proteome ^24^ and the absence of other members of the SKI complex in this protein fraction, the ATP requirement for vacuolar RNA transport, and the *ski2* phenotypes described here, support the hypothesis that SKI2 is a component of a plant RNautophagy-like mechanism. Alternatively, an indirect effect could be invoked. The cytoplasmic RNA exosome and the SKI complex have an important role in the degradation of mRNA decay intermediates. If 5’ fragments produced by endonucleolytic cleavage of abundant transcripts are not removed by the cytoplasmic RNA exosome, they can be recognized as aberrant transcripts by the cellular post-transcriptional gene silencing machinery, triggering silencing of the corresponding mRNA ^49–51^. Plants with knock-down of core components of the exosome can have pleiotropic phenotypes, such as altered cuticular wax composition, due to the silencing of genes associated with these pathways ^52,53^. Thus, it is possible that the *ski2* mutant is missing a key component of the RNA transport system due to defects in RNA exosome function suppressing silencing of endogenous transcripts.

In conclusion, plant vacuoles have the ability to actively import RNA through a mechanism that is functionally analogous to RNautophagy, but mechanistically distinct. The RNA helicase SKI2 is needed for a fully active mechanism and it is likely to be a component of the transport system. This novel mechanism can translocate rRNA and thus may have a significant role in RNA salvage and nucleotide homeostasis in plant cells, and its ability to transport other RNAs, represented by random sequence short oligonucleotides in our assay, could indicate that this mechanism is also important in the turnover of other cytoplasmic RNA species. The assays developed for this work should facilitate the molecular dissection of this new mechanism.

## Materials and Methods

### Plant growth and *Arabidopsis thaliana* genotypes

*Arabidopsis thaliana* plants used for vacuole analyses were grown in soil under short day conditions (10hr light/14hr dark) at 22°C for 6 weeks. For whole seedling microscopy, seeds were surface-sterilized in 33% (v/v) bleach, 1% Triton X-100 for 20min followed by cold treatment for ≥2 days. Plants were grown for seven days under long day conditions (16hr light/8hr dark) at 22°C on nutrient solid half-strength Murashige-Skoog medium with vitamins (MSP09; Caisson Labs), 1% sucrose, 2.4mM MES pH5.7, and 0.8% (w/v) phytoagar (PTP01; Caisson Labs) as described ^54^. Arabidopsis Columbia-0 accession was used as wild type control. T-DNA insertion mutants used in this study were *rns2-2* ^6^, *ski2-5* (SALK_141579C), *rh40-1* (CS824414) and *rh40-2* (SALK_054998). To generate the *ski2-5*/*SKI2* line, The *SKI2* cDNA sequence was amplified from wild type cDNA using CloneAmp HiFi PCR Premix (Takara) and cloned into the pPZP212 binary vector, resulting in a SKI2 gene driven by 35S promoter at the 5’ end, and with a MYC tag sequence at the 3’ end of the insert. Transgenic plants were generated by the floral dip method. Genotyping was carried out using standard qPCR and PCR techniques. Mutants and transgenic lines are available from the corresponding authors upon request. A list of primers used in this work is provided in Supplementary Table 1.

### qRT-PCR-based assay of in vitro RNA transport into vacuoles

Vacuoles were purified from *rns2-2* plants and membrane integrity confirmed by neutral red staining and brightfield microscopy according to Robert et al. ^55^. Two vacuole preparations were used for each experiment. Purified vacuoles were combined into one tube and gently homogenized by pipetting with cut tips. A 200μl aliquot of vacuole preparation was added to each reaction tube. 5μl of 2U/μl apyrase (A6410; Sigma Aldrich) was added to minus ATP samples and incubated for 5 minutes at room temperature to remove residual ATP. 5μg *Drosophila melanogaster* total RNA [purified as previously described ^56^] were dissolved in nuclease free water, then added to the reaction along with 18mM adenosine 5’-triphosphate (ATP) magnesium salt (A9187; Sigma Aldrich) for plus ATP samples. Following gentle mixing with cut pipet tips, samples were incubated at room temperature for 5 minutes to allow transport. Samples were then incubated with 200μg RNaseA (19101; Qiagen) for 20min on ice to remove residual Drosophila RNA outside the vacuole. 10μl Recombinant RNasein© Ribonuclease inhibitor (N2515; Promega) was added to inhibit RNaseA activity and samples were flash frozen in liquid nitrogen until analyzed. Upon thawing for analysis, 18mM ATP was added to samples without prior ATP addition, as the ATP concentration affects acid phosphatase activity. Vacuoles were homogenized by vortexing and spun down prior to aliquoting 35μl for use in acid phosphatase analysis as described ^7^. RNA was extracted from the remaining vacuoles using a RNeasy Plant Mini Kit (74904; Qiagen). For this, vacuoles were combined with 600µl of lysis buffer RLT (Qiagen) and RNA extracted according to the manufacturer’s protocol. Samples were DNase-treated using on column DNaseI (79254; Qiagen) and RNA eluted with 30μl of water. A 13μl aliquot of extracted vacuolar RNA was used for cDNA synthesis by a qScript Flex cDNA kit (95049; Quanta BioSciences) using random primers with a final cDNA dilution volume of 80μl. The quality of cDNA was tested by semi-quantitative PCR. qRT-PCR was carried out using PerfeCta SYBR Green Fast Mix, Low ROX (95074; Quanta BioSciences) in a Stratagene Mx4000 multiplex quantitative PCR system (Agilent Technologies) using primers for Drosophila 18S and 28S rRNA. qPCR results were normalized to the vacuole marker enzyme acid phosphatase activity ^57,58^. Samples not containing vacuoles were assigned an acid phosphatase value based on each experiment’s average acid phosphatase activity for normalization purposes.

### Visualizing RNA transport into the vacuole using Alexa Fluor 488 labeled RNA

Vacuoles were purified from wild type and mutant (*rns2-2*, *ski2-5*, *rh40-1, rh40-2*) rosette leaves. 200μl purified vacuoles were used for each sample following gentle sample homogenization by pipetting, and kept on ice. Vacuoles were treated with apyrase and 10mM ATP. 1μl of 413pmol/μl Alexa488-RNA was added. The Alexa488-RNA was a custom machine mixed random 15 base polyribonucleotide sequence with an Alexa488 molecule conjugated to the 3’ end of the oligo and HPLC purified (Integrated DNA technologies). As a negative control, Alexa488-RNA was incubated with 50μg of RNaseA (19101; Qiagen) for 20-30 minutes at room temperature in darkness to remove RNA from the Alexa488 fluorophore. Prior to visualization by fluorescence microscopy the samples were diluted 10-fold with ice-cold vacuole buffer ^55^ to dilute residual Alexa488 fluorophore. Vacuoles were then imaged by fluorescence or confocal microscopy. For inhibitor studies, 200μl purified wild-type vacuoles were prepared as above. Vacuoles were treated for 5 minutes on ice with either 10mM adenosine 5’-(β,γ-imido) triphosphate lithium salt hydrate (A2647; Sigma Aldrich), 15µM gramicidin (G5002; Sigma Aldrich), or sodium orthovanadate (S6508; Sigma Aldrich). Following incubation with either orthovanadate or gramicidin, 10mM ATP and 1μl of 413pmol/μl Alexa488-RNA was added to samples to measure vacuole RNA transport. Samples were incubated on ice for 20mins, diluted and visualized by confocal or fluorescence microscopy.

### Microscopy for RNA transport and autophagy quantification

Vacuoles were imaged by confocal microscopy using a NikonC1si confocal scanning system attached to a 90i microscope (Nikon Instruments). For quantification experiments, vacuoles were visualized using a Zeiss Axio Imager.A2 upright fluorescent microscope (Carl Zeiss Inc.) with a 10x objective and green fluorescent protein filter. Exposure levels were set using ATP treated samples and all subsequent images were taken with the same settings. Vacuoles were classified by relative intensity (2= high intensity, 1= low intensity, 0= no intensity), and fluorescence intensity of each vacuole was quantified using Fiji software ^18^. A circular region within the vacuole being measured was quantified, and average intensities for each sample were statistically compared.

Transgenic *ski2-5* plants expressing GFP-ATG8e were generated as described ^59^ and GFP-ATG8e-labeled autophagosomes were visualized by confocal microscopy and counted per unit area of root. All experiments were done in triplicate per genotype.

### Immunoblotting

ATG8 protein accumulation was analyzed as described previously, with minor modifications ^10^. Briefly, seven-day old WT, *ski2-5, rh40-1, rh40-2, ski2-5/SKI2* and *rns2-2* seedlings were incubated with or without 50 μM inosine (final concentration) in liquid ½ MS medium supplemented or not with 1μM final concentration of concanamycin A (ConcA) for 12 hours, and total protein was separated by SDS–PAGE followed by immunoblotting with anti-ATG8a antibody (AS14 2769, Agrisera).

### Statistical analysis

Except as indicated, all data were analyzed in triplicate using R (version 4.0.3). Two-tailed Student’s *t*-tests were computed for pairwise comparison of means of data sets, with the statistical significance level set to 0.05. One-way analysis of variance (ANOVA) was followed with Tukey’s multiple comparisons test or post-hoc pairwise *t*-test.

## Abbreviations

ConcA: concanamycin A
FL-RNA: fluorescent RNA
LAMP2C: Lysosome-associated membrane glycoprotein 2
RH40: RNA Helicase 40
RNS2: ribonuclease2
SIDT2: SID1 transmembrane family member 2
SKI2: superkiller2

## Acknowledgements

This work was supported by a grant from the National Science Foundation (MCB-1714996) to G.C.M. and D.C.B.

## Contributions

B.E.F., G.C.M. and D.C.B. conceived the project and designed experiments. B.E.F., M.M.W., A-Y.L., S.C.M., Y.M., Z.K, V.R. and X.L. performed experiments and analyzed data. B.E.F., G.C.M. and D.C.B. prepared the manuscript. All authors revised the manuscript.

## Supplementary Figures

**Figure S1.**
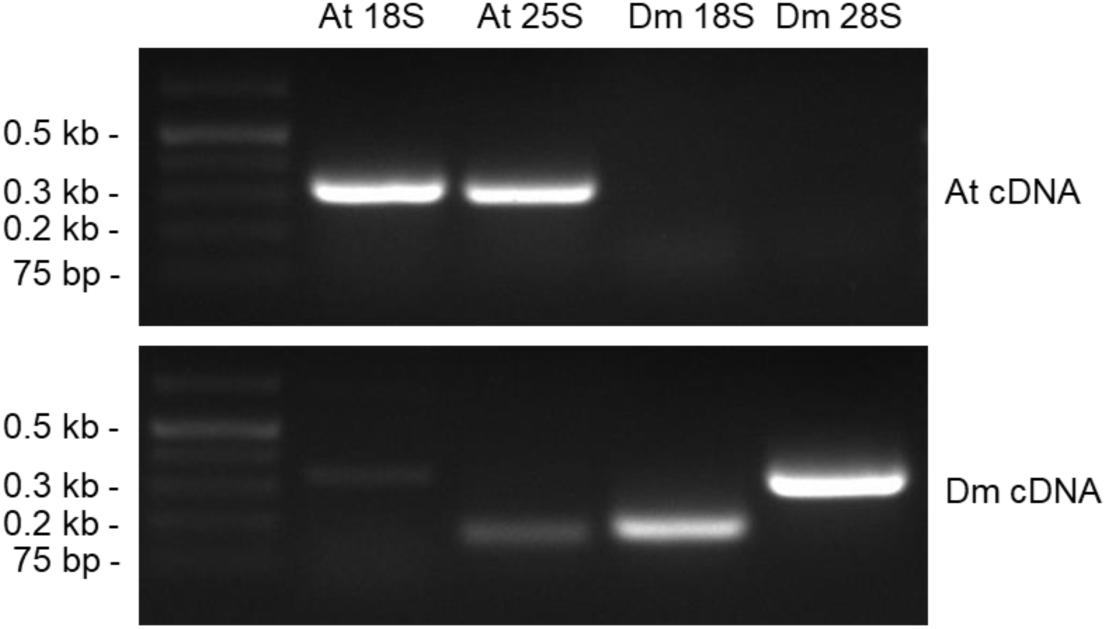
*Drosophila melanogaster* rRNA-specific primers. Total RNA was extracted from Arabidopsis rosette leaves and Drosophila ovaries and tested by RT-PCR using primers against either Drosophila or Arabidopsis rRNA sequences. The Drosophila primers are specific for Drosophila 18S and 28S rRNA. The positions of size markers are shown at left. At: *Arabidopsis thaliana*, Dm: *Drosophila melanogaster*.

**Figure S2.**
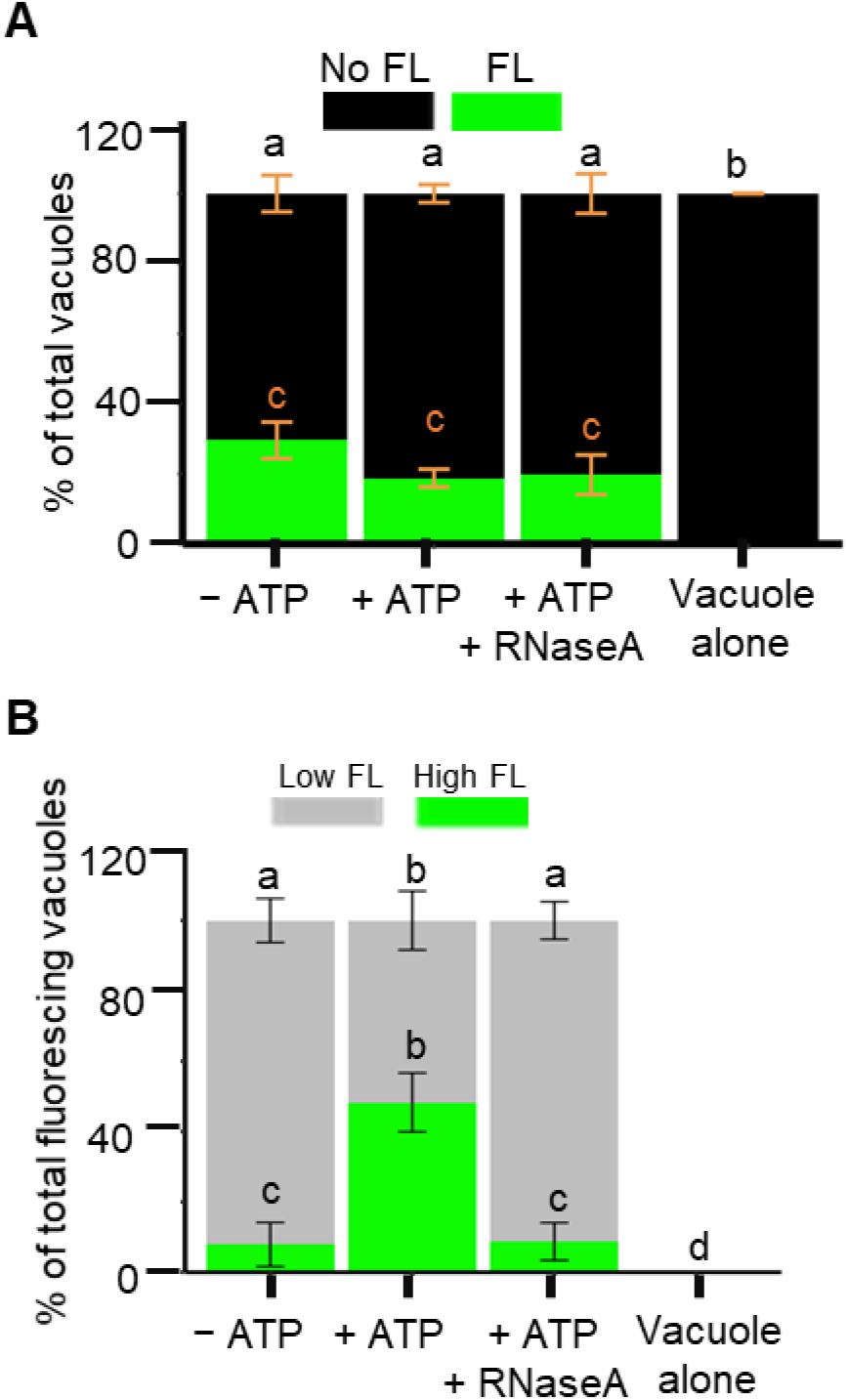
Alexa488-tagged-RNA is transported into *rns2-2* vacuoles in vitro. (**A**) The number of *rns2-2* vacuoles at zero, low, and high fluorescence intensity for three biological replicates was quantified and normalized to the total number of vacuoles. (**B**) Fraction of low and high fluorescence vacuoles normalized to total number of fluorescent vacuoles. Error bars represent ± SD. Similar letters indicate no significant difference according to one-way analysis of variance (ANOVA) with Tukey’s multiple comparisons test; For (A), a, b; b, c; a, c; p < 0.0001; For (B), a, b; b, c; b, e; p < 0.01, a, c; a, e; p < 0.0001, c, e; p < 0.05. FL, fluorescence.

**Figure S3.**
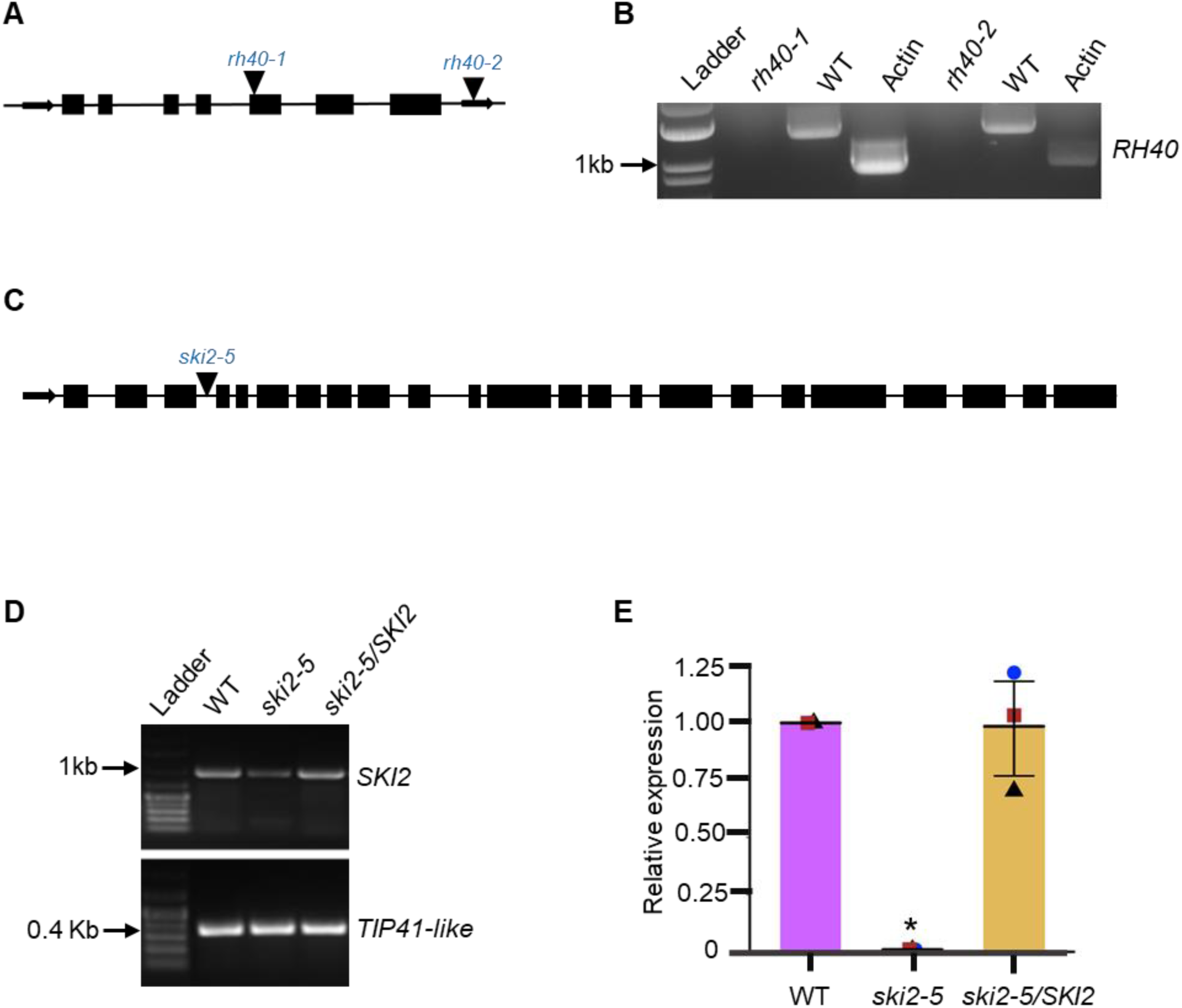
Characterization of the *rh40-1*, *rh40-2* and *ski2-5* mutant lines. (**A**) Schematic of the two independent T-DNA insertion alleles of *rh40*. (**B**) RT-PCR was used to detect the presence of *RH40* transcripts. The lack of *RH40* transcripts in the mutant lines indicates that *rh40-1* and *rh40-2* are null mutants. (**C**) Schematic of the T-DNA insertion allele of *ski2-5*. (**D**) RT-PCR was used to detect the presence of *SKI2* transcripts in the *ski2-5* mutants. A reduced amount of transcript was observed in the mutant line. (**E**). Q-PCR was used to quantify the relative amount of *SKI2* transcript present in the *ski2-5* mutant. Results indicate that *SKI2* is expressed in the *ski2-5* mutant at less than 1% of the wild type level. The *ski2-5/SKI2* line corresponds to the *ski2-5* mutant complemented with a wild type *SKI2* gene, and shows expression similar to wild type. Error bars indicate ± standard deviation from three biological replicates, with means for each replicate represented by the blue circle, black triangle and brown square. Asterisk indicates significant difference according to pairwise Student’s two-sided *t*-test; *p* < 0.001.

**Table S1.**
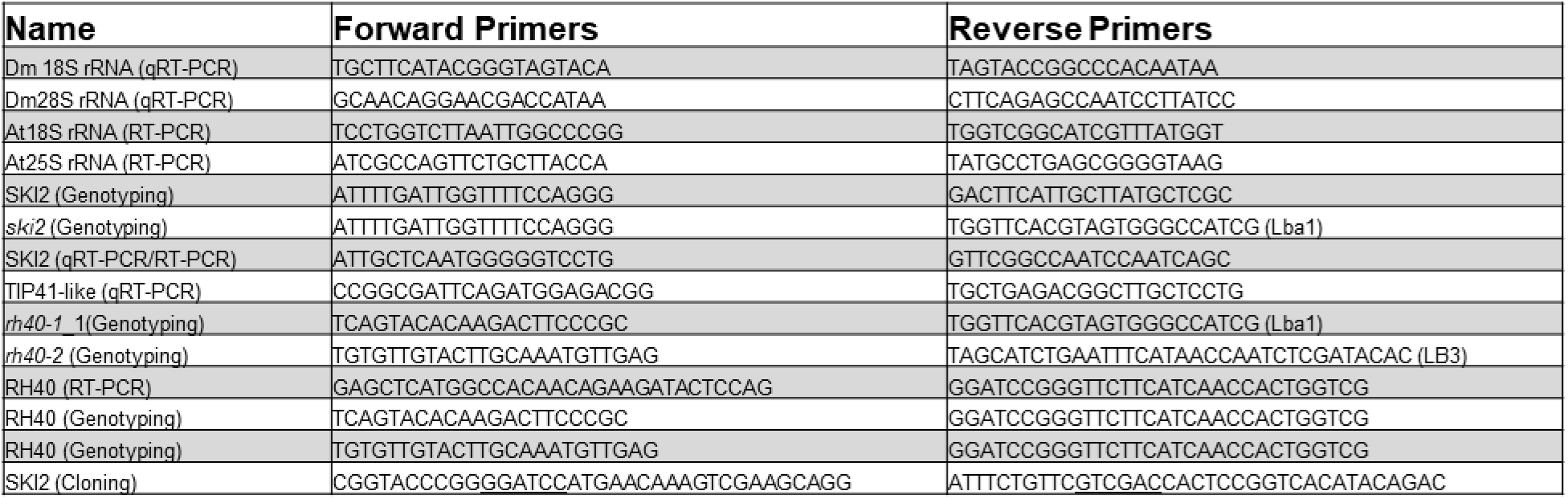
Primers used in this study.

